# Dengue virus preferentially uses human and mosquito non-optimal codons

**DOI:** 10.1101/2023.06.14.544804

**Authors:** Luciana A Castellano, Ryan J McNamara, Horacio M Pallares, Andrea V Gamarnik, Diego E Alvarez, Ariel A Bazzini

## Abstract

Codon optimality refers to the effect codon composition has on messenger RNA (mRNA) stability and translation level and implies that synonymous codons are not silent from a regulatory point of view. Here, we investigated the adaptation of virus genomes to the host optimality code using mosquito-borne dengue virus (DENV) as a model. We defined which codons are associated with unstable and stable (non-optimal and optimal codons, respectively) mRNAs in mosquito cells and showed that DENV preferentially uses non-optimal codons and avoids codons that are defined as optimal in either human or mosquito cells. Human genes enriched in the codons preferentially and frequently used by DENV are up-regulated during infection, and so is the tRNA decoding the non-optimal and DENV preferentially used codon for arginine. We found that synonymous mutations towards DENV’s preferred non-optimal codons (e.g., AGA) increase fitness of DENV during serial passaging in human or mosquito cells. Finally, our analyses revealed that hundreds of viruses preferentially use non-optimal codons, with those infecting a single host displaying an even stronger bias, suggesting that synonymous codon choice is a key aspect of host-pathogen interaction.

## Introduction

Dengue is the most prevalent mosquito-borne viral disease in the world, causing an estimated 390 million infections each year^1^. This disease is caused by four (DENV1-4) distinct but genetically related serotypes. DENV is a single-stranded positive-sense RNA virus^2^ that is spread to humans through the bite of infected *Aedes* species mosquitoes. Given the limited coding capacity of their RNA genome, RNA viruses rely on the host translation machinery to synthesize their protein components. Hence, the viral genome is thought to be under selective pressure to adapt to the host cell translation resources^3^.

Previous work has revealed that vertebrate viruses have an AT-rich genome and that their codon usage does not correlate with that of their hosts^4, 5^. In addition to codon usage (i.e. codon frequency), ‘codon optimality’ is an important post-transcriptional mechanism that shapes gene expression based on the codon composition, impacting homeostatic mRNA and protein levels from yeast to human cells^6–16^. ‘Codon optimality’ refers to the effects that specific codons have on mRNA stability and translation efficiency^6–9, 17^. Codons that enhance mRNA stability and translation efficiency are defined as ‘optimal codons’, while ‘non-optimal’ codons have the opposite effect (Figure Supplement 1A)^8^. Thus, mRNAs enriched in optimal codons tend to be more stable and more efficiently translated than mRNAs enriched in non-optimal codons. While the molecular mechanisms involved in codon optimality remain poorly characterized, optimal codons are associated with higher levels of tRNA and higher charged to uncharged tRNA ratios^6, 8, 9, 11, 18, 19^. This codon-mediated regulation suggests that while synonymous codons encode the same amino acids, they are not silent from a regulatory point of view and can impact mRNA and protein abundance^8, 9, 15, 20^.

As DENV is an RNA virus which alternates between human and mosquito hosts, it has evolved unique strategies to overcome the barriers imposed by switching between vertebrate and invertebrate hosts^21–24^, representing an interesting model to interrogate the relationship between viral codon choice and host codon optimality. Here, we defined the codon optimality code in mosquito cells and investigated DENV codon preference relative to both host codon optimality codes. We also investigated whether host gene expression, including tRNA abundance, changes in a codon-dependent manner upon DENV infection and whether synonymous mutations are beneficial for the virus. This work revealed that DENV and many other viruses preferentially use non-optimal codons compared to their host which has important implications for understanding host-pathogen interactions.

## Results

### Many viruses preferentially use non-optimal codons relative to humans

To determine which codons DENV preferentially uses relative to its host, a Relative Synonymous Codon Usage (RSCU) fold change was calculated as the ratio of RSCU observed in DENV to that of its host (e.g. human) per amino acid. RSCU is independent of amino acid frequency (Figure 1A and 1B). Then, the correlation between the RSCU fold change and the codon stability coefficient (CSC) from human cells^8^, a measure of codon optimality, was calculated to investigate the relationship between DENV’s codon preference and the hosts’ codon optimality. A significant negative correlation was observed between the RSCU fold change of DENV2 and codon optimality (R = -0.26, *p* = 0.045) (Figure 1C), suggesting that DENV preferentially uses non-optimal codons relative to humans. For example, DENV preferentially uses AGA to encode arginine relative to humans; AGA is the most non-optimal codon encoding arginine in human cells (Figure 1D). A similar trend was observed for other amino acids, such as isoleucine and glutamine (Figure 1D).

**Figure 1:**
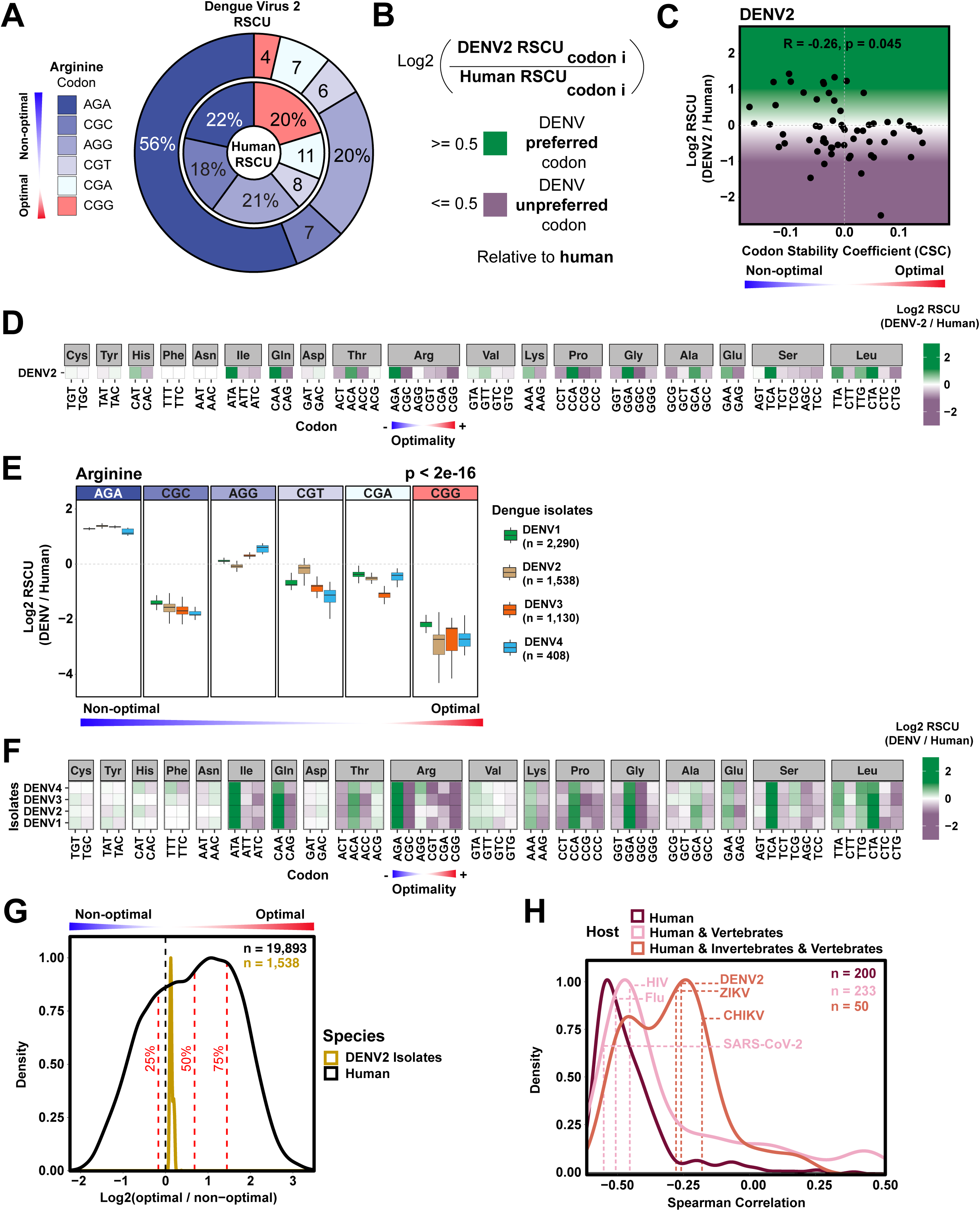
Several RNA viruses preferentially use non-optimal codons relative to humans. A) Donut chart showing the percentage of dengue virus (DENV) Relative Synonymous Codon Usage (RSCU) and human RSCU for arginine codons. Codons are labeled based on human optimality (red = optimal, blue = non-optimal). B) The RSCU fold change was used as a metric for DENV’s codon preference relative to human and it was calculated as the log2 ratio of RSCU observed in DENV compared to human per synonymous codon per amino acid. Codons showing a RSCU fold change greater than 0.5 are DENV preferred codons (green), codons with a RSCU fold change less than 0.5 are DENV unpreferred codons (purple). C) Scatterplot showing the RSCU fold change (relative to human) for DENV2 and human codon stability coefficient (CSC). R = -0.26, *p* = 0.045, Spearman rank correlation. D) Heatmap showing the RSCU fold change (relative to human) for DENV2. Codons are ordered in increasing human optimality within each amino acid. E) Boxplot showing the RSCU fold change (relative to human) of the synonymous codons encoding arginine for 5,366 DENV isolates spanning the four serotypes (DENV1-4). Human CSC indicated by color of codon (red = optimal, blue = non-optimal). *p* < 2e-16, ANOVA, RSCU fold change relative to human ∼ codon. F) Heatmap showing the RSCU fold change (relative to human) for 5,366 DENV isolates spanning the four DENV serotypes (DENV1-4). Codons are ordered in increasing human optimality within each amino acid. G) Density plot showing log2 ratio of optimal to non-optimal codon frequency for human endogenous genes and DENV2 isolates. Quantiles for human density indicated by red dashed lines. *p* = 1.59e-98. H) Density plot showing Spearman rank correlation between RSCU fold change (relative to human) and human CSC for human-infecting viruses. Viruses are split based on their host. Spearman rank correlation of particular viruses are indicated with labeled dashed lines.

To investigate whether this preference exists in the genomes of DENV natural isolates, the RSCU fold change was calculated from 5,366 world-wide isolated DENV sequences spanning the four serotypes. The DENV isolates’ relative preference for arginine codons is not equally distributed (p < 2e-16) (Figure 1E). The most non-optimal codon encoding arginine, AGA, is the most preferred codon by all four serotypes (RSCU FC: 1.32 ± 0.07) (Figure 1E). The strong preference for AGA might suggest evolutionary pressure to select this non-optimal codon in the genome of DENV isolates. Moreover, each of the serotypes exhibits a preference for the most or second most non-optimal codon for several other amino acids: isoleucine, glutamine, threonine, arginine, lysine, proline, glycine, glutamic acid and serine (Figure 1F). These results indicate that this trend in preference for non-optimal codons is conserved across DENV serotypes, despite the fact the serotypes only share ∼70-75% of their amino acid and nucleotide sequences (Figure Supplement 1B). The global codon optimality of DENV2 isolates falls in the 33.9 ± 0.01 percentile of human gene optimality, indicating that natural circulating strains of DENV2 viruses preferentially use non-optimal codons relative to humans (*p* = 1.59e-98) (Figure 1G).

To ask whether the preference for non-optimal codons was a general trend across viruses and not a specific observation for DENV, we calculated the correlation between the RSCU fold change and human CSC for 483 viruses that infect humans. The vast majority exhibit a similar preference for non-optimal codons (negative correlation), including influenza virus, HIV and SARS-CoV-2 (Figure 1H). Viruses that only infect humans or humans and vertebrates, presented a stronger negative correlation than viruses that infect humans and invertebrates, suggesting that host dependent evolutionary pressure may shape the codon preference (Figure 1H).

### Mosquito cells exhibit codon optimality

To investigate whether the relationship between the codon preference of DENV and codon optimality is host dependent, since DENV infects humans and mosquitoes, we determined whether codon optimality exists in *Ae. albopictus* C6/36 cells, using methods we and others have applied in other systems^6, 8, 9, 13, 14, 16^. After identifying the codon optimality properties in mosquito (optimal or non-optimal) (Figure 2), we investigated the relationship between the viral codon preference and codon optimality in each host (human and mosquito) (Figure 3).

**Figure 2:**
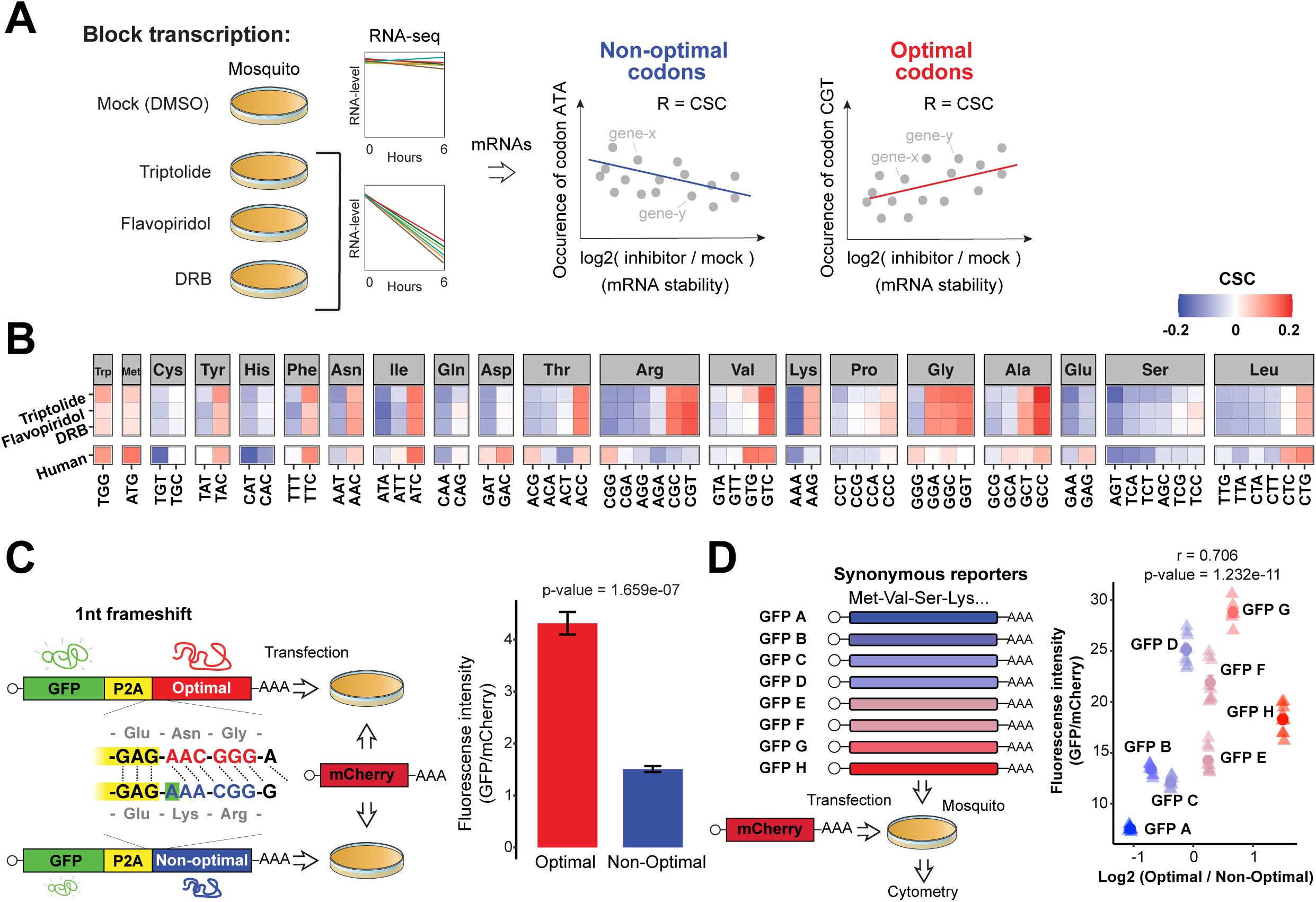
Mosquito C6/36 cells exhibit codon optimality. A) Diagram illustrating endogenous mRNA decay profiles approach to determine the codon optimality code in mosquito C6/36 cells. RNA-seq was performed at 6 hours after mock treated (DMSO) or blocking transcription with Flavopiridol, Triptolide or 5,6-dichloro-1-beta-D-ribofuranosylbenzimidazole (DRB). The codon stabilization coefficient (CSC) was calculated as the Pearson correlation coefficient between the occurrence of each codon and the log2 ratio of mRNA level between treated and mock cells (metric for mRNA stability). B) Heatmap showing the CSC calculated in human cells and in mosquito cells using the indicated transcription inhibitors to measure mRNA stability. C) Schematic of the 1nt-out of frame reporters: two mRNAs that differ in codon composition due to a single nucleotide insertion (A in blue, highlighted in green), causing a frameshift. The encoding GFP fluorescent protein is followed by a cis-acting hydrolase element (P2A) and then by the coding region enriched in optimal or non-optimal codons due to the frame shift. P2A causes ribosome skipping, thus the GFP is not fused to the optimal or non-optimal encoded proteins. These reporters were co-transfected into *Ae. albopictus* C6/36 cells with a vector encoding for mCherry as an internal control. Bar plot showing that the 1nt-out of frame reporter enriched in optimal codons displayed higher GFP/mCherry fluorescense intensity than its non-optimal counterpart measured by flow cytometry analysis. *p* = 1.659e-7, unpaired t-test. D) Illustration of 8 GFP encoding mRNAs differing only in synonymous mutations. All GFP variants were co-transfected into *Ae. albopictus* C6/36 cells with a vector encoding for mCherry as an internal control. Scatter plot showing a positive correlation between the log2 ratio of optimal to non-optimal codon frequency for all these GFP variants (based on mosquito optimality) and GFP fluorescence intensity in mosquito C6/36 transfected cells measured by flow cytometry analysis. R = 0.706, *p* = 1.232e-11, Spearman rank correlation.

**Figure 3:**
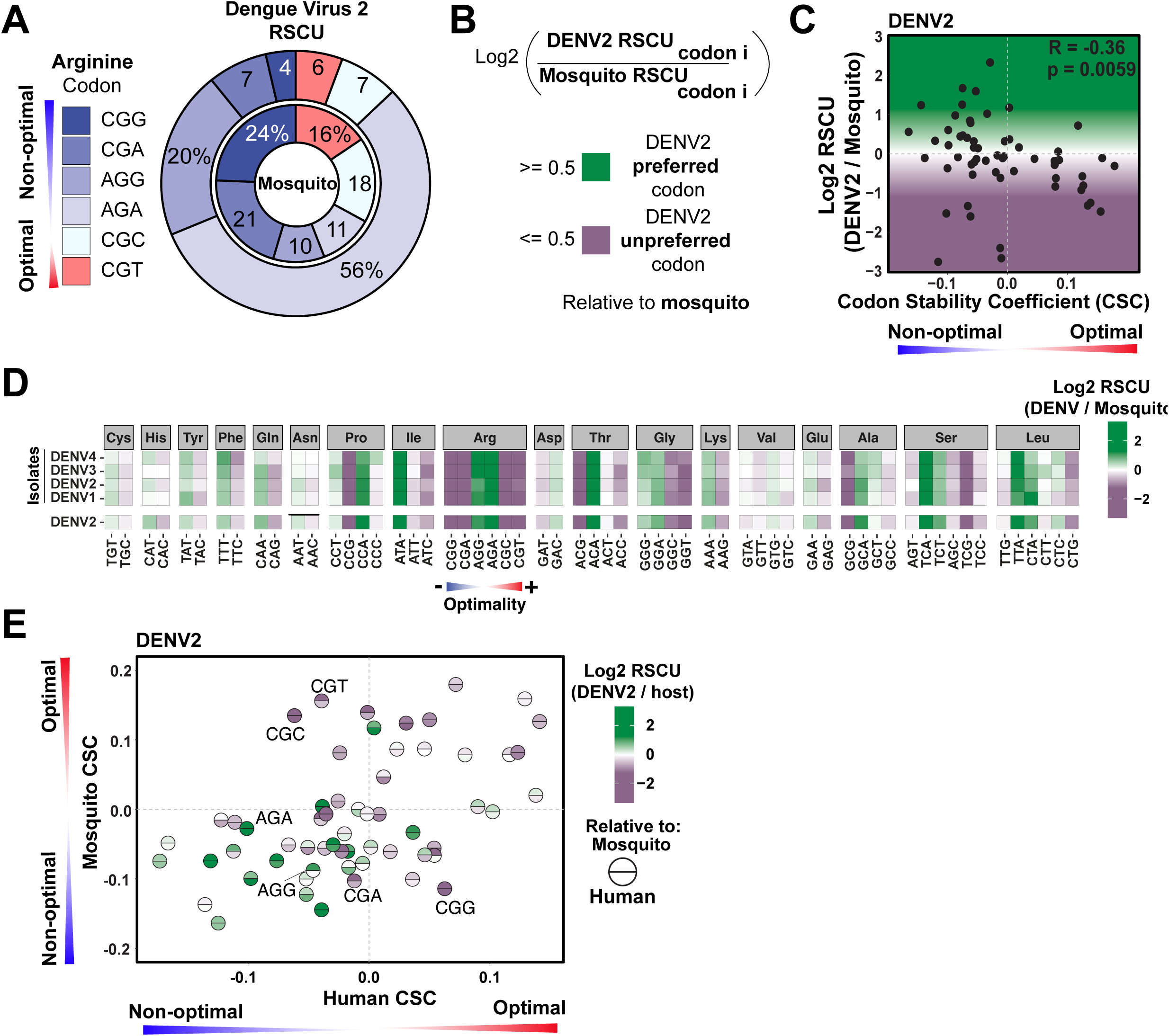
Dengue virus preferentially uses non-optimal codons relative to mosquitoes. A) Donut chart showing the percentage of dengue virus (DENV) Relative Synonymous Codon Usage (RSCU) and mosquito RSCU for arginine codons. Codons are labeled based on mosquito optimality (red = optimal, blue = non-optimal). B) The RSCU fold change was used as a metric for DENV’s codon preference relative to mosquito and it was calculated as the log2 ratio of RSCU observed in DENV compared to mosquito per synonymous codon per amino acid. Codons showing a RSCU fold change greater than 0.5 are DENV preferred codons (green), codons with a RSCU fold change less than 0.5 are DENV unpreferred codons (purple). C) Scatterplot showing the RSCU fold change (relative to mosquito) for DENV2 strain 16684 and mosquito codon stability coefficient (CSC). R = -0.36, *p* = 0.0059, Spearman rank correlation. D) Heatmap showing the RSCU fold change (relative to mosquito) for 5,366 DENV isolates spanning the four DENV serotypes (DENV1-4) and DENV2 strain 16684. Codons are ordered in increasing mosquito optimality within each amino acid. E) Scatterplot showing mosquito and human CSC. Each circle represents a codon. Color of the circle indicates DENV2’s RSCU fold change relative to mosquito (top half) or human (bottom half). Preferred codons in both hosts (top and bottom green) cluster in the bottom left quadrant (non-optimal in both hosts). R = 0.46, *p* = 0.00032, Spearman rank correlation.

To determine the potential regulatory properties of the codons affecting mRNA stability and protein expression (optimal or non-optimal codons) in mosquitoes, *Ae. albopictus* C6/36 cells were treated with DMSO or with one of three different transcription inhibitors: Flavopiridol, Triptolide, and 5,6-dichloro-1-beta-D-ribofuranosylbenzimidazole (DRB) (Figure 2A). RNA-Seq was performed at 6 hours post-treatment. The fold change of the mRNA level between transcription inhibitor and DMSO treatments was calculated as a metric for mRNA stability. For each codon, the codon stability coefficient (CSC) was calculated as the Pearson correlation between mRNA stability and codon occurrence^9^. Codons exhibiting a positive correlation are referred to as optimal codons, while codons exhibiting a negative correlation are referred to as non-optimal codons (Figure 2A).

The CSCs calculated using the three transcription inhibitors showed significant correlation (DRB-Flavopiridol: R = 0.993, *p* = 3.886e-58; DRB-Triptolide: R = 0.966, *p* = 6.803e-37; Triptolide-Flavopiridol: R = 0.952, *p* = 2.606e-58), indicating reproducibility independent of the transcription inhibitor used (Figure 2B and Figure Supplement 2A). Hence, the codon optimality code of *Ae. albopictus* C6/36 cells was determined based on the combined results for the three independent transcription inhibitors used in the assays (Figure 2B). The mosquito codon optimality code showed stronger correlation with Drosophila^16^ (R = 0.788, p < 2.2e-16), than zebrafish^6^ (R = 0.503, *p* = 4.591e-5) or human^8^ (R = 0.477, *p* = 1.218e-4) (Figure Supplement 2B). While there are certain amino acids, such as valine, for which the codon optimality in mosquito and human are similar, there are other amino acids with different codon optimality properties. For example, CGG is the most optimal codon encoding arginine in humans, but it is the most non-optimal in mosquito. Conversely, CGT is the most optimal codon encoding arginine in mosquito, but it is non-optimal in human. AGA is non-optimal in both species (Figure 2B).

To validate that the regulatory information is encrypted in a codon-dependent manner and not simply in the nucleotide composition, we used a pair of reporters that differed by a single nucleotide insertion (1nt frameshift). The extra nucleotide causes a frameshift that converts a ‘non-optimal’ sequence (enriched in non-optimal codons) into an ‘optimal’ coding sequence (enriched in optimal codons), otherwise keeping the nucleotide composition nearly identical (Figure 2C). The coding sequence of green fluorescent protein (GFP) was followed by a ribosome skipping sequence (P2A)^25, 26^ and a coding region enriched in either optimal or non-optimal codons (due to 1 nucleotide frameshift) (Figure 2C). The reporter enriched in optimal codons showed higher level of GFP compared to the counterpart reporter (enriched in non-optimal codons) in mosquito cells co-transfected with mCherry as an internal control (*p* = 1.659e-7, unpaired t-test) (Figure 2C). Moreover, as the 1nt-out of frame reporters encode different amino acids, we also compared the expression of GFP reporters which differ only in synonymous codons^20^ (Figure 2D). We observed a positive correlation between the percentage of optimal and non-optimal codons (defined in mosquito) and the fluorescence intensity in mosquito cells transfected with each of the GFP variants (R = 0.706, *p* = 1.232e-11, Spearman correlation) (Figure 2D). All together these results indicate that the codons contain regulatory properties and confirm that codon optimality exists in mosquito cells.

### Dengue virus preferentially uses codons that are not optimal in human or mosquitoes

After defining the codon optimality code of *Ae. albopictus*, we investigated the relationship between synonymous codon preferences of DENV and codon optimality in mosquitoes. First, to determine which codons DENV preferentially uses relative to mosquito, the RSCU fold change was calculated as the ratio of RSCU observed in DENV to that of mosquito per amino acid (Figure 3A and 3B). A significant negative correlation was observed between the RSCU fold change of DENV2 relative to mosquito and mosquito codon optimality (R = -0.36, *p* = 0.0059) (Figure 3C). Specifically, for amino acids such as isoleucine, threonine, glycine, alanine, serine and leucine, the most preferred codon used by the four DENV serotypes (shown in green) relative to mosquito tends to be non-optimal (Figure 3D). In the case of arginine, the codon AGA is less non-optimal in mosquitoes than in humans, but it is still non-optimal (Figure 2B) and strongly preferred by all DENV serotypes relative to mosquito (Figure 3D). Moreover, the thousands of worldwide DENV isolates showed the preference for AGA relative to mosquito (Figure 3D & Figure Supplement 3A). Overall, our findings suggest that DENV prefers to use codons that are non-optimal in mosquitoes, consistent with our results in humans (Figure 1).

To further explore the relationship between relative codon preference and host optimality, the codon optimality of mosquitoes was plotted against the codon optimality of humans (Figure 3E). The top half of each circle (codon) was color-coded based on the RSCU fold-change of DENV2 relative to mosquito and the bottom half relative to human. Codons preferentially used by DENV relative to both human and mosquito (green top and green bottom) cluster in the lower-left quadrant. This indicates that DENV2 prefers codons that are non-optimal in both hosts (Figure 3E), while codons that are optimal in one or both hosts tend to not be preferentially used by DENV. In sum, DENV does not preferentially use codons that are defined as optimal in either of its hosts (human and/or mosquito).

### Human genes enriched in the codons used by dengue virus are upregulated upon dengue virus infection

Our results indicate DENV prefers non-optimal codons relative to humans and mosquitoes. To investigate whether changes in host gene expression upon DENV infection are regulated in a codon-dependent manner, we analyzed single-cell sequencing data of DENV2 infected human Huh7 cells^27^. First, we defined a set of ‘denguenized’ codons as the codons that are both preferentially used by DENV relative to humans (more than 0.5 log2 RSCU fold change) and are one of the 16 most frequently used codons by DENV. Conversely, we called ‘not-denguenized’ codons as a group of codons that are not preferentially used by DENV relative to human (less than -0.5 log2 RSCU fold change) and are one of the 16 least frequently used codons by DENV (Figure 4A and Figure Supplement 4A and 4B).

**Figure 4:**
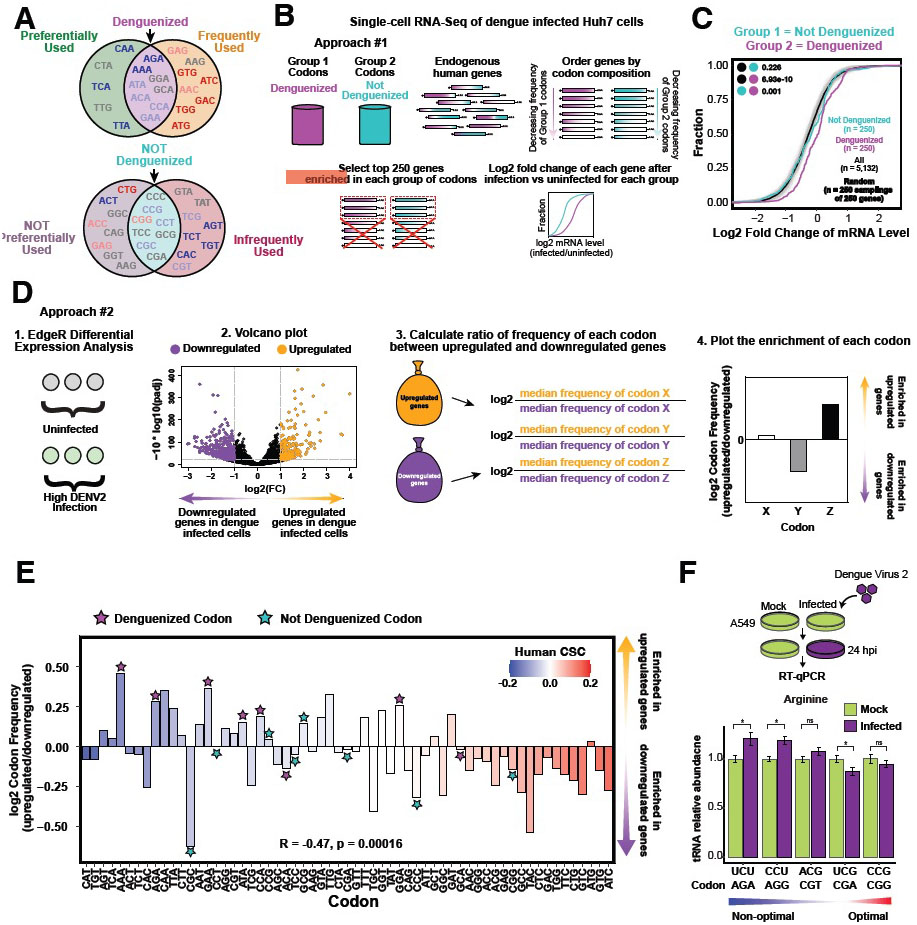
Codon-mediated regulation of gene expression in dengue infected human cells. A) Venn diagrams showing the codons that are preferentially or not preferentially used by dengue virus (DENV) relative to human, and the ones that are frequently or infrequently used in DENV2’s genome. Codons that are preferentially and frequently used by DENV were called ‘denguenized’ and the codons not preferentially and infrequently used were called ‘not denguenized’. Color of the codon indicates human CSC (red = optimal, blue = non-optimal, grey = neutral). B) Diagram of the mRNA level analysis of genes enriched in ‘denguenized’ and ‘not denguenized’ codons (Approach #1). Human endogenous genes enriched in each codon group were identified and their mRNA level fold change upon DENV2 infection was calculated from single cell sequencing of infected Huh7 cells using edgeR package. C) Cumulative distribution of mRNA level fold change from human endogenous mRNAs enriched in ‘denguenized’ and ‘not denguenized’ codons, upon DENV2 infection. All other genes are shown in black. 250 samplings of 250 random endogenous genes from ‘All’ group shown in light gray. *p*-values indicated, Wilcoxon rank-sum test. D) Schematic of the codon frequency analysis of genes upregulated/downregulated upon DENV2 infection of Huh7 cells (Approach #2). Cells were ordered in increasing DENV expression and differential expression analysis was performed on uninfected and high-infection groups using edgeR package. Codon frequency of upregulated and downregulated genes was calculated and plotted as the log2 ratio of the median frequency of each codon in the group of upregulated and downregulated genes. E) Barplot showing enrichment of each codon in the upregulated/downregulated genes. Codons with a value greater than 0 are enriched in the upregulated group, codons with a value less than 0 are enriched in the downregulated group. Codons are ordered in increasing human CSC (red bars = optimal, blue bars = non-optimal, white bars = neutral). R = -0.47, *p* = 0.00016, Spearman rank correlation. Stars indicate ‘denguenized’ (purple) and ‘not denguenized’ (teal) codons. Non-optimal, ‘denguenized’ codons are enriched in upregulated genes (*p* = 0.000136, unpaired Wilcoxon test). F) qRT-PCR analysis showing the relative quantification of arginine tRNA levels in mock infected and DENV infected human A549 cells. Results are shown as the averages of tRNA abundance relative to tRNAHisGTG ± standard error of the mean from two independent experiments with three biological replicates per experiment (\**p* < 0.05, unpaired t-test).

Two independent approaches were taken to investigate whether there is a codon-meditated effect on host gene expression upon DENV infection. First, two groups of genes were created by selecting the 250 human endogenous genes most enriched in ‘denguenized’ or ‘not-denguenized’ codons. The mRNA level fold change between infected and uninfected cells was then calculated for each group of genes (Figure 4B). Genes enriched in ‘denguenized’ codons were upregulated upon infection compared to genes enriched in ‘not-denguenized’ codons (*p* = 1e-3, Wilcoxon rank-sum test) (Figure 4C).

In the second approach, the differentially expressed genes (up-and down-regulated) between infected vs. uninfected cells were selected (adjusted *p*-value < 0.01, and fold change > 1 or < -1, respectively), and then the codon composition of each group was compared (Figure 4D). The group of up-regulated genes during DENV infection were enriched in non-optimal codons and depleted in optimal codons, relative to the downregulated genes (R = -0.47, *p* = 0.00016, Spearman correlation) (Figure 4E). The group of up-regulated genes during infection also showed a higher enrichment of the ‘denguenized’ codons relative to the group of down-regulated genes (*p* = 0.000136, unpaired Wilcoxon test) (Figure 4E).

As tRNA level correlates with the regulatory properties of the codons (optimal or non-optimal)^6, 8, 9^, we hypothesized that these codon-dependent changes in gene expression may be explained by changes in the tRNA pool upon DENV infection. Interestingly, the relative level of arginine tRNA decoding AGA (non-optimal, ‘denguenized’) and AGG (non-optimal) were upregulated in A549 human cells infected with DENV compared to mock infected cells (Figure 4F, *p* < 0.05, unpaired t-test). In contrast, the relative level of arginine tRNA decoding CGA (neutral, ‘not-denguenized’) was down-regulated in infected cells compared to mock infected cells (Figure 4F, *p* < 0.05, unpaired t-test). Non-significant changes in the level of arginine tRNA decoding CGT (slightly non-optimal) and CCG (optimal) codons were observed between the infected and uninfected cells (Figure 4F, *p* > 0.05, unpaired t-test). Together these results support the idea that DENV infection alters tRNA availability and may underlie the codon-based changes observed in human gene expression during infection.

### Dengue virus synonymous mutations towards dengue virus’ preferred codons tend to increase the viral relative fitness during adaptation to human or mosquito

Following the rationale that synonymous mutations are not silent from a regulatory point of view, and that DENV preferentially uses non-optimal codons, we investigated the adaptive effect of synonymous mutations by analyzing sequencing data of DENV in *in-vitro* directed evolution through serial passaging in a single host cell line^28^. Analyses were focused on synonymous mutations arising after nine serial passages following the initial transfection of DENV2 strain 16681 viral RNA into human Huh7 or *Aedes albopictus* C6/36 mosquito cells (Figure 5A). The relative fitness associated with all possible mutations (beneficial, deleterious, neutral, or lethal) across the DENV genome was calculated based on the frequency trajectory of a given allele over time relative to its mutation rate^28^. To evaluate the fitness effect of synonymous substitutions, we defined the Mean Relative Fitness (MRF) for each codon as the average across the relative fitness of every possible synonymous mutation to the codon across the DENV genome (Figure 5B). The MRF of a codon represents the average relative fitness associated with a synonymous mutation to the codon, independent of the position in the genome where the mutation occurs. Therefore, an MRF of 1 indicates a neutral mutation, on average. A mutation with an MRF greater than 1 is considered beneficial on average, indicating that a synonymous mutation towards the particular codon increases viral fitness. Conversely, an MRF less than 1 suggests a deleterious mutation on average which decreases viral fitness.

**Figure 5:**
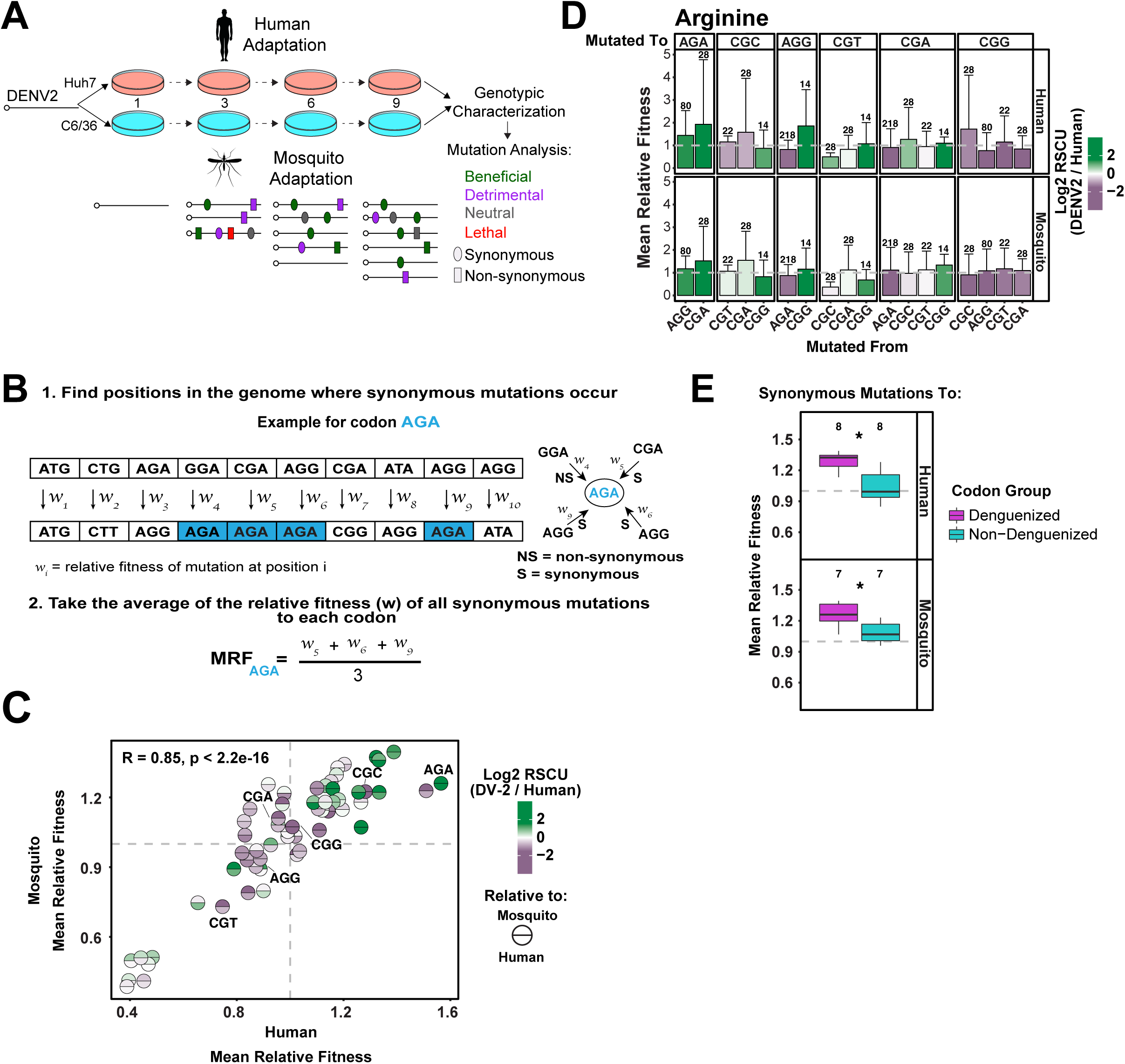
Relative fitness of dengue virus mutations upon infection correlates with dengue virus’ codon preference relative to mosquitoes and humans. A) Outline of *in vitro* dengue virus (DENV) evolution experiment performed by Dolan et al. DENV RNA (Serotype 2/16881/Thailand/1985) was electroporated into mosquito (C6/36) or human (Huh7) cell lines, and the resulting viral stocks were passaged for nine passages in biological duplicates. After passage, samples of virus from each passaged population were subject to genotypic characterization by ultra-deep sequencing using the CirSeq procedure. B) Diagram of calculation of the Mean Relative Fitness (MRF) of synonymous substitutions in the DENV genome during adaptation to human or mosquito cells. First, for each codon, we selected all synonymous mutations *to* that codon. Second, we used sequencing data from the 9^th^ passage to identify the positions in the DENV genome where these mutations occurred and obtain the relative fitness (w) associated with each synonymous substitution at each position. Finally, the MRF of each codon was calculated by taking the average across the relative fitnesses of every possible synonymous mutation *to* the codon across the DENV genome. The MRF represents the average relative fitness associated with synonymous mutations *to* a codon independent of the position in the genome where the mutation occurred. C) Scatter plot showing the Mean Relative Fitness (MRF) after adaptation to mosquito or human cells. Each codon, represented by a circle, was labeled with DENV2’s codon preference (RSCU fold change) relative to mosquito (top half) and human (bottom half) hosts. An MRF greater than 1.0 indicates the mutation towards that particular codon increases the viral fitness, on average. DENV’s preferred codons relative to both hosts (top and bottom green) cluster in the top right quadrant, suggesting synonymous mutations towards these codons increase the fitness of DENV during adaptation to mosquito and human cells. R = 0.85, *p* < 2.2e-16, Spearman rank correlation. D) Bar plot comparing the Mean Relative Fitness (MRF) of synonymous mutations within arginine during adaptation to human (top) or mosquito (bottom) cells. Bars are labeled with DENV2’s codon preference (RSCU fold change) relative to human (top) or mosquito (bottom) hosts. An MRF greater than 1.0 indicates the mutation towards that particular codon increases the viral fitness, on average. E) Boxplot showing the Mean Relative Fitness (MRF) of synonymous mutations to the group of ‘denguenized’ (purple) and ‘not denguenized’ (teal) codons during adaptation to human (top) or mosquito (bottom) cells. Synonymous mutations towards ‘denguenized’ codons showed higher MRF than mutations towards ‘not denguenized’ codons, suggesting that the former are more beneficial for viral fitness compared to the latter.

Interestingly, the MRF in mosquito and human are correlated (R = 0.85, *p* < 2.2e-16) and the codons preferentially used by DENV relative to both mosquito and human (green top and green bottom) tend to have positive MRF (Figure 5C). For example, within the codons encoding arginine, synonymous mutations going from AGG or CGA to AGA showed MRF greater than 1, while synonymous mutations from AGA to AGG or CGA showed MRF less than 1, suggesting that it is beneficial for the virus to select AGA and detrimental to lose AGA (Figure 5D). Similar results were obtained by analyzing all the ‘denguenized’ and ‘not denguenized’ codons (Figure 5E). Specifically, synonymous mutations towards ‘denguenized’ codons showed higher MRF than mutations towards ‘non-denguenized’ codons, suggesting that the former are more beneficial for viral fitness compared to the latter (Figure 5E). Our results indicate that adaptation of DENV to human or mosquito cells results in the selection of synonymous mutations towards DENV’s preferred codons, which tend to be non-optimal, suggesting that DENV has evolved a preferred codon usage away from host codon optimality.

## Discussion

In this study, we provide diverse experimental and analytical evidence that indicate that viral synonymous codon choice is an important aspect of host-pathogen co-evolution. Several general conclusions arise from this work. First, DENV preferentially uses non-optimal codons and avoids codons that are defined as optimal in either human or mosquito cells (Figure 3 and Figure 6). Second, similar codon preference was observed in the four DENV serotypes, including thousands of world-wide isolates, despite the fact they share less than 75% amino-acid similarity (Figure 1). These points are clearly illustrated by codons encoding arginine, as AGA, the most non-optimal codon in human, was not only the most preferentially used by DENV, but also shows the lowest dispersion of RSCU within the thousands of world-wide isolates suggesting an evolutionary pressure to select this codon (Figure 1). Third, host genes (humans) enriched in the codons preferentially and frequently used by DENV, defined as ‘denguenized’ are up-regulated during DENV infection, suggesting a co-evolution between the codons used by the virus and by the host genes induced upon infection (Figure 4 and Figure 6). The fact that the tRNA decoding the most non-optimal and preferentially used codon encoding for arginine is up-regulated upon infection; and, that the tRNA decoding the most optimal and not preferred codons by DENV are down-regulated or not changing, suggests that the regulatory properties (optimal or non-optimal) of the codons might change during infection (Figure 4 and Figure 6).

**Figure 6:**
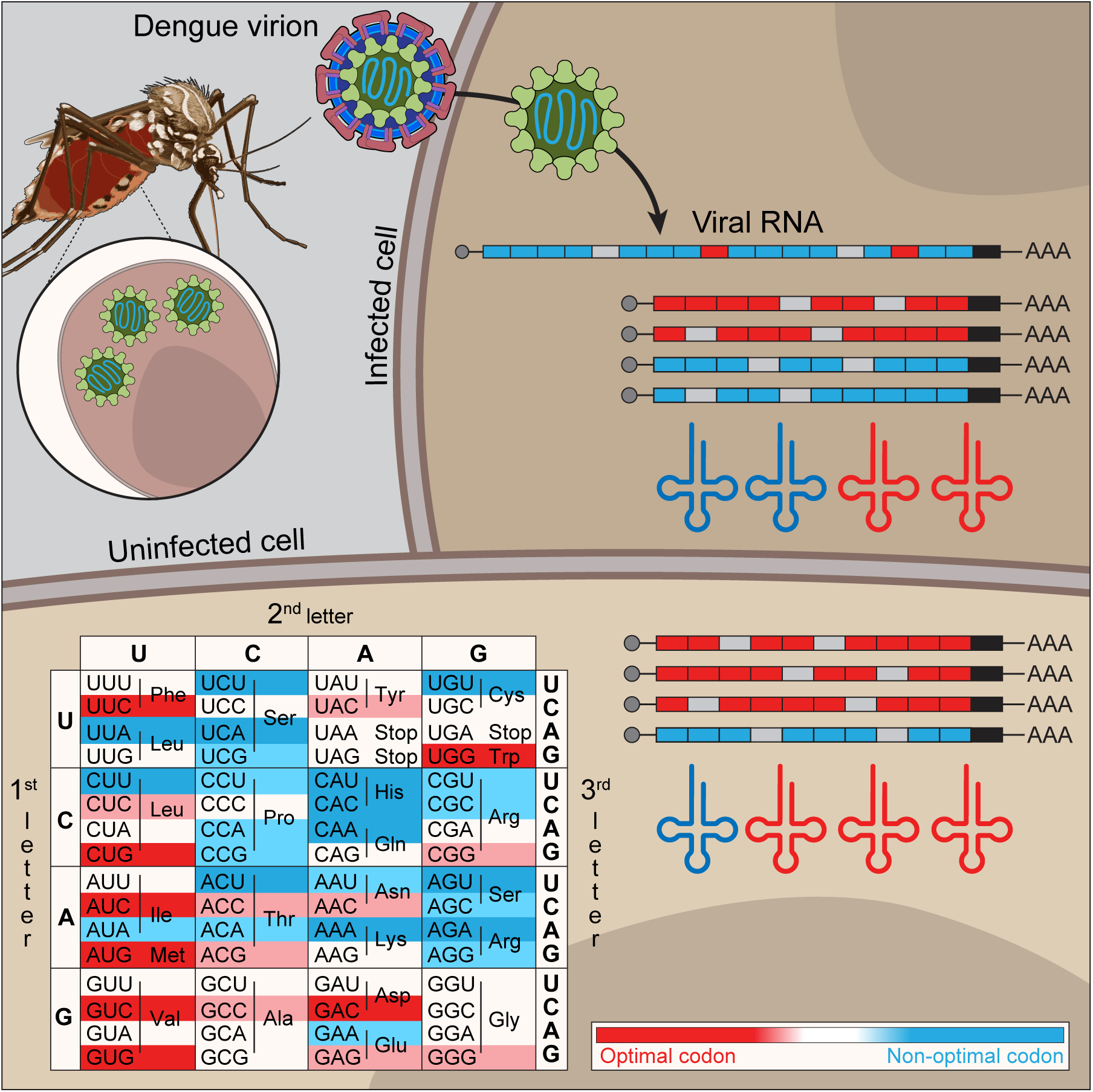
Model of codon-dependent regulation during dengue virus infection. Dengue virus preferentially uses non-optimal codons relative to both human and mosquito hosts. Upon infection, human genes enriched in non-optimal codons are upregulated as well as arginine tRNA decoding the most non-optimal and dengue preferred codon.

A fourth general observation is that analysis of synonymous mutations after host-restricted serial passages in human or mosquito cells showed that synonymous mutations towards the preferred non-optimal codons (e.g., AGA) increase DENV fitness during infection. Our analysis of the correlation between codon optimality and codon preference showed that many viruses, including DENV, tend to use non-optimal codons as defined by their host and thus, DENV is not an exception. These correlations were stronger for viruses that only infect humans or humans and vertebrates, compared to viruses that infect humans and invertebrates, suggesting that codon preference is under evolutionary pressure and depends on the evolutionary distance of their host (Figure 1).

Several hypotheses on virus-host interactions emerge from these observations. For example, viruses might avoid optimal codons because increasing viral RNA stability and translation efficiency might be detrimental, triggering an early antiviral response and/or causing the lysis of the host cell. Our findings are in line with evidence that suggests that viruses with codon usage too similar to that of their host could be harmful to host cells as a result of the high level of expression of viral genes and consequent tRNA depletion effect^4^. Additionally, codon composition influences RNA structure, translation elongation, and protein folding^8, 9, 29–31^, so we cannot rule out the possibility that codon-mediated effects on these might benefit the virus. In order to miss-regulate host genes and favor the cellular environment towards the virus, viruses might affect tRNAs that are normally expressed at low levels and service non-optimal codons. tRNA encoding for optimal codons are highly expressed relative to their demand and so we hypothesize that it would be more challenging to affect gene expression through misregulation of optimal codons^6, 8, 18^. tRNA can be regulated at multiple levels. For example, viruses such as HIV^32^, polyomavirus (SV40)^33^, adenovirus^34, 35^, MHV68^36^, Epstein Barr Virus^37^ and HSV-1^38^, stimulate Pol III transcription. Additionally, changes in tRNA availability in HSV-1 infected cells were linked with interferon activation^39^. The changes in tRNA level have been proposed to favor translation of viral proteins^32, 40^. Beyond tRNA levels, tRNA modifications play a major role in tRNA function, including stability and translation fidelity^41, 42^. For instance, it has been proposed that there is a codon-specific reprogramming of translation via tRNA modification in chikungunya (CHIKV) infected human cells^43^. Interestingly, AGA (Arg), GAA (Glu), AAA (Lys), CAA (Gln), and GGA (Gly) codons are preferred by both CHIKV and DENV, and they are targets of the KIAA1456 enzyme involved in the tRNA modifications that favor decoding of these A-ending codons over the G-ending ones^43^. Moreover, tRNA-derived fragments (tRFs) can have regulatory function, and, upon respiratory syncytial virus (RSV) infection, tRFs promote viral replication^44^. Therefore, it is plausible that changes in tRNA levels and tRNA modifications, as well as an upregulation of tRF upon infection, might play a role in optimizing the host environment to favor translation of the viral RNA^45^.

There are other cellular contexts where tRNAs change with a concomitated change in the expression the genes enriched in those codons. For example, the tRNA repertoire varies depending on the differentiation or proliferation status of the cells and is tightly coordinated with changes in the transcriptome^46^. Up-regulation of specific tRNAs in human breast cancer cells promotes breast cancer metastasis through a remodeling of protein expression by enhancing stability and/or ribosome occupancy of transcripts enriched for their cognate codons^47^. Arginine limitation in colorectal cancer cells represses arginine tRNAs, leading to ribosomal stalling at arginine codons and a proteomic shift to arginine low proteins^48^. Moreover, a higher abundance of m^7^G-modified tRNA Arg-TCT has been reported to drive oncogenic transformation through reshaping gene expression by enhancing stability and translation efficiency of mRNAs enriched in the corresponding AGA codon^49^. Loss of function of one central nerve specific isodecoder of tRNA Arg-TCT induces ribosome stalling at AGA codons causing neurodegeneration in mice^50^.

Our work raises three questions for future study: What is special about the tRNA decoding AGA, which systematically appears to be the most regulable and which is related to neurodegenerative disease, cancer, and now virus infection? Are the changes in the tRNA pool initiated upon DENV infection beneficial for the virus and/or for the host? Mechanistically, how is the tRNA availability being modulated upon viral infection? It would be interesting to know whether modulations of the tRNA pool are actively induced by the virus or by the host in response to viral infection. In summary, this work showing that DENV and other viruses preferentially use non-optimal codons compared to their host has important implications for understanding the evolution of host-pathogen interactions.

## Acknowledgements

Authors thank Dr Robb Krumlauf and Sara Bancroft (Stowers Institute) for suggestions and critical reading of the manuscript, and Mark Miller for designing the graphical abstract. Authors also thank the following Stowers Core facilities: Cells, Tissues and Organoids Center, Media Prep, Sequencing and Discovery Genomics, Cytometry, Automation and PCR Technology, and Computational Biology. Authors are thankful to members of the Bazzini laboratory for discussions, especially Gabriel da Silva Pescador, Anthony Treichel and Dr Qiushuang Wu for technical support. This study was supported by the Stowers Institute for Medical Research.

A.A.B and D.E.A were awarded a Pew Innovation Fund and A.A B. with the US National Institutes of Health (NIH-R01 GM136849 and NIH R21OD034161). This work was performed as part of thesis research for L.A.C, Graduate School of the Stowers Institute for Medical Research. The funders had no role in study design, data collection and interpretation, or the decision to submit the work for publication.

## Author contribution

DEA, and AAB designed and conceived the project. LAC, RJM and AAB interpreted the data. LAC conceived, planned, and performed the experiments and data analysis. RJM performed data curation and data analysis. HMP provided support for experiments. AVG provided reagents and materials. AAB supervised the work. LAC, RJM, DEA and AAB wrote the manuscript with input from the other authors. The authors read and approved the final manuscript.

---

**Figure supplement 1.**
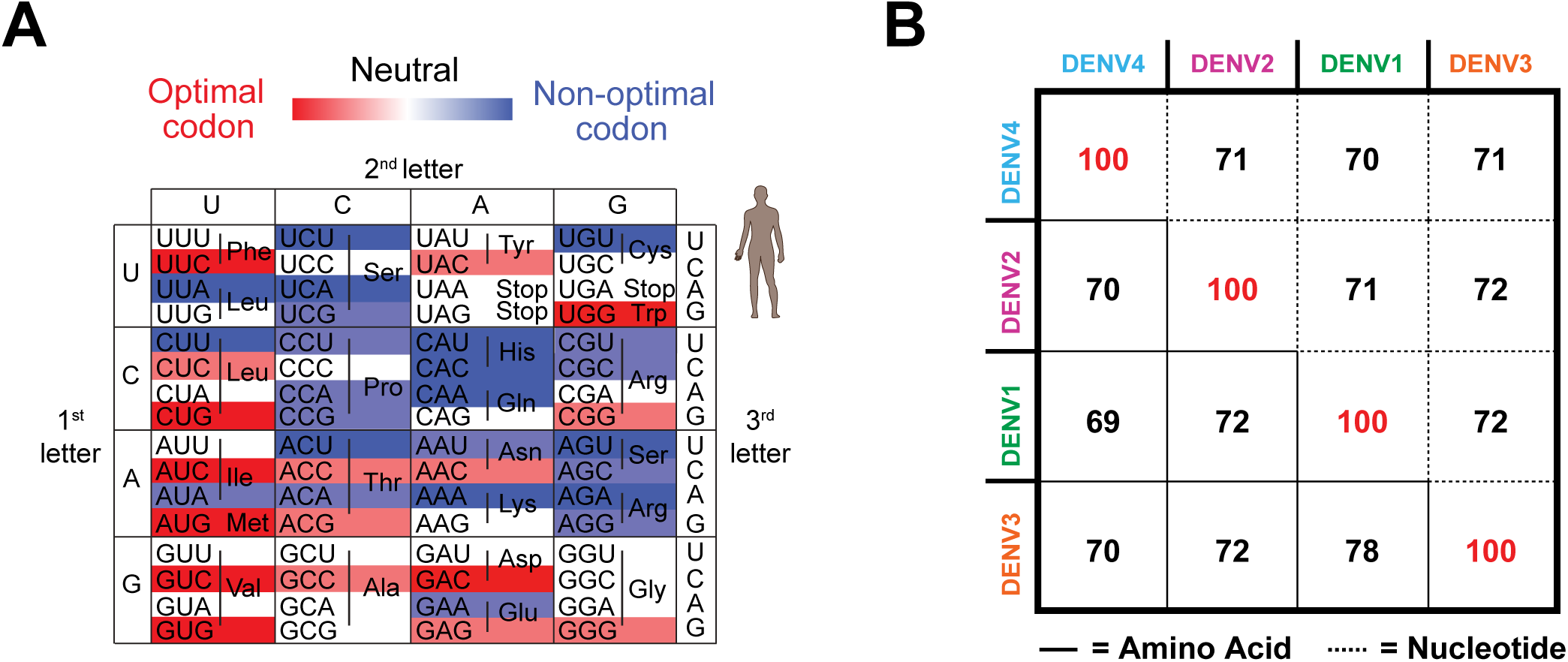
A) Heatmap showing human CSC. Optimal codons highlighted in red, non-optimal codons highlighted in blue. B) Matrix showing the four DENV serotypes’ (DENV 1-4) similarity. Lower triangle indicates amino acid similarity, upper triangle indicates nucleotide similarity.

**Figure supplement 2.**
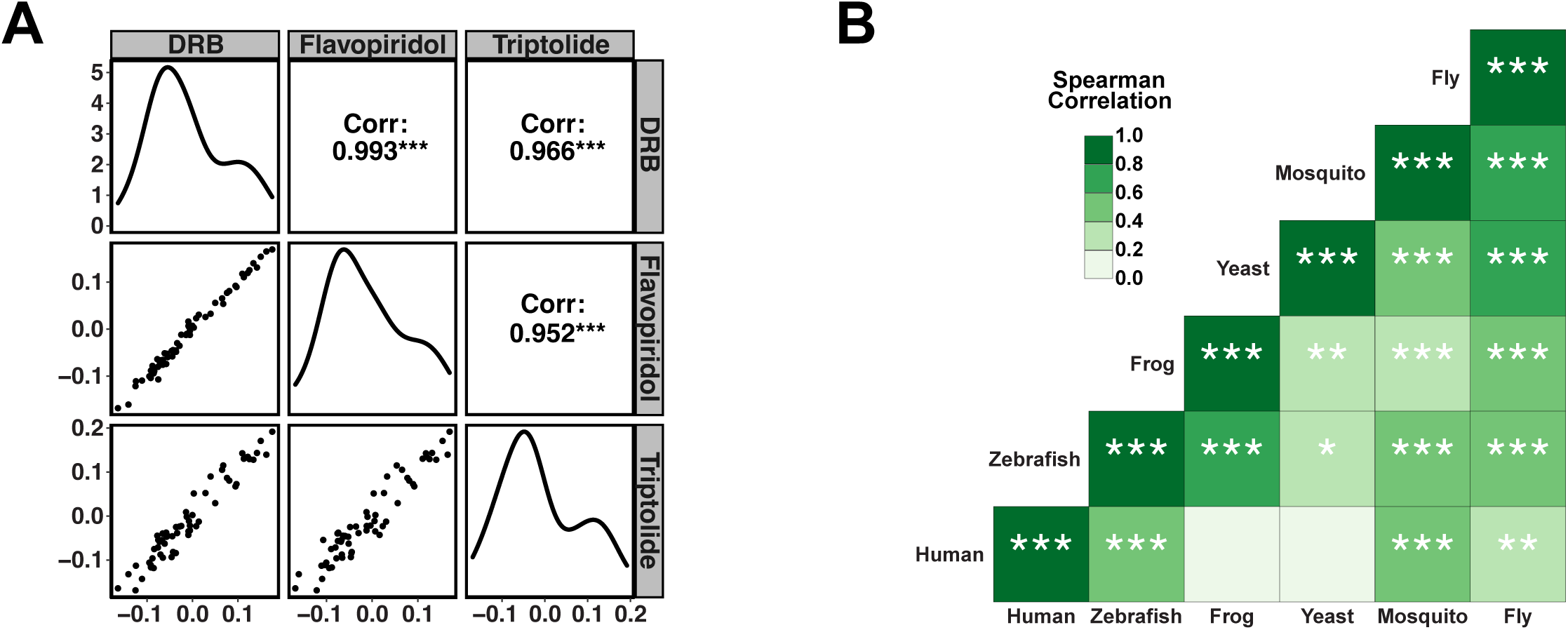
A) Pair plot showing the CSC calculated in mosquito C6/36 cells using indicated transcription inhibitors. Lower triangle shows scatterplots of CSCs between inhibitors. Diagonal shows density plot of CSC for each inhibitor. Upper triangle shows Pearson correlation coefficient of CSCs between inhibitors. B) Heatmap showing Spearman rank correlations between known CSCs in indicated species. Color of the tile indicates correlation coefficient. \**p* < 0.05, \*\**p* < 0.01, \*\*\**p* < 0.001.

**Figure supplement 3.**
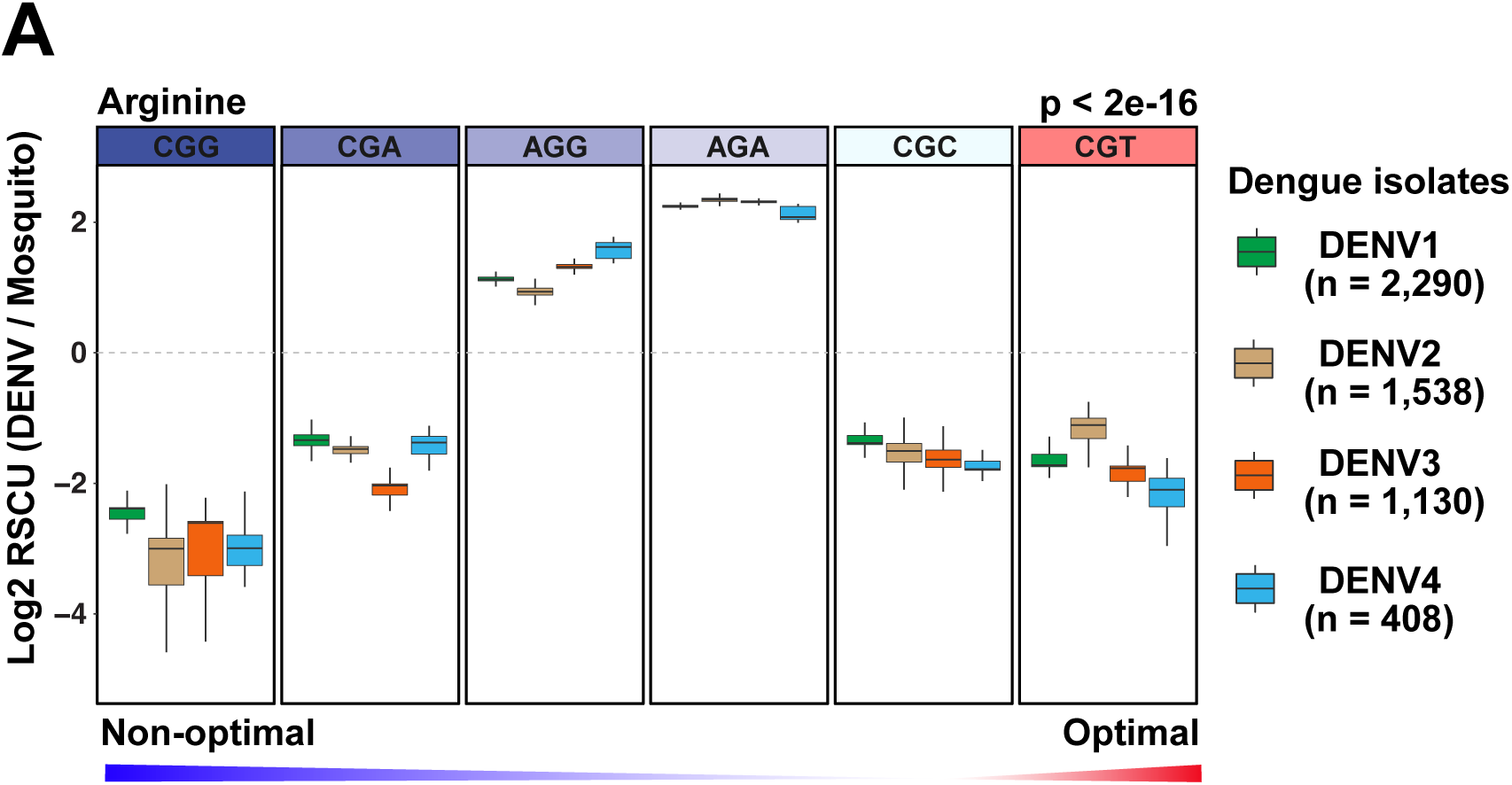
A) Boxplot showing the RSCU fold change (relative to mosquito) of the synonymous codons encoding arginine for 5,366 DENV isolates spanning the four serotypes (DENV1-4). Mosquito CSC indicated by color of codon (red = optimal, blue = non-optimal). *p* < 2e-16, ANOVA, RSCU fold change relative to mosquito ∼ codon.

**Figure supplement 4.**
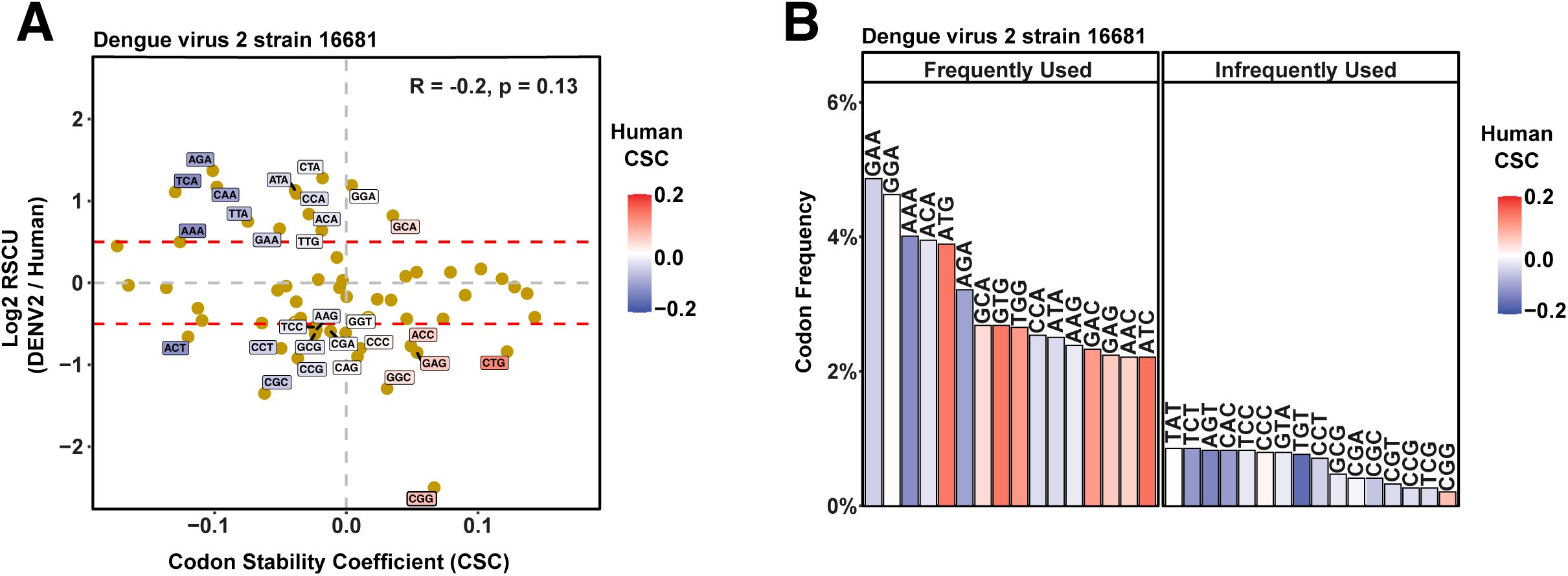
A) Scatterplot showing the RSCU fold change (relative to human) for DENV2 strain 16681 and human codon stability coefficient (CSC). Labels indicate ‘preferentially used’ (log2(RSCU fold change relative to human) ≥ 0.5) and ‘not preferentially used’ (log2(RSCU fold change relative to human) ≤ -0.5). R = -0.2, *p* = 0.13, Spearman rank correlation. B) Barplot showing the frequency of the 16 most used (‘frequently used’) and the 16 least used (‘infrequently used’) codons in the DENV2 strain 16681 genome.

